# Dissociation between biomechanical stiffness and sEMG activity in trapezius and lumbar skeletal muscles in steady postures

**DOI:** 10.1101/2024.06.27.600991

**Authors:** Alar Veraksitš, Märt Reinvee, Jaan Ereline, Helena Gapeyeva, Tatjana Kums, Georg Gavronski, Mati Pääsuke, Eero Vasar

**Affiliations:** Chair of Physiology, Institute of Biomedicine and Translational Medicine, University of Tartu, Estonia; Institute of Technology, Estonian University of Life Sciences, Tartu, Estonia; Institute of Sport Sciences and Physiotherapy, University of Tartu, Estonia

**Keywords:** Electromyography, muscle stiffness, standing, lying, sitting

## Abstract

**Background:** Prolonged sitting posture and sedentary behaviour, spent mostly in sitting are harmful for general health. The low back and shoulder area are the most vulnerable. In these regions the sEMG registration of neuromuscular activity shows low activity in steady postures. Stiffness of according muscle can be measured by myotonometry. We were not able to find any direct comparison between these parameters although separately, direct correlations between contraction force and sEMG or stiffness have been clearly established.

**Research question:** Whether and how stiffness is modulated by neuromuscular activity in standing, lying or in different sitting postures in these vulnerable regions.

**Methods:** The muscle’s biomechanical stiffness (measured with MyotonPRO) and mean power frequency (MPF) with amplitude (AMP) on surface electromyography (sEMG) were registered in the upper part of *musculus trapezius* (UT) and *musculus erector spinae* (ES, at the level of L4 vertebrae). Nine healthy physically active males aged 19–46 (mean±SD, 28.6±10.9 years), participated in the study. The standing, prone, and three sitting postures where studied. The latter were distinguished by the back-tight-angle (BTA): 1) sitting on a common chair, straight back, BTA 90°), 2) slumped sitting on the same chair (BTA<90°), and 3) sitting on an experimental chair with a convex base, BTA 115-120°.

**Results and Significance:** Muscle stiffness did not correlate with either of the sEMG parameters in ES but did so only in low grade with the AMP in UT (Spearman rank ρ=0.24, p=0.02). It was interesting that contrary to UT, in ES a significant positive correlation (ρ=0.24, p=0.02) was noted between MPF and AMP. It is likely that the steady body position under the Earth’s g-force may be ensured by the biomechanical characteristics of the tissue rather than neuromuscular activity. This can be explained by incompressible nature of soft tissues and be a less resource-consuming strategy.

**Highlights:**

- Muscle stiffness in the low back is similar to standing straight and lying.
- In steady postures muscle stiffness does not correlate with neuromuscular activity.
- Body position is consolidated by the muscle’s biomechanical stiffness.

## 1. Introduction

If the body posture is stable, the shoulder and low back areas show no considerable amount of neuromuscular activity. This has been observed in the lumbar area during several stable postures and also during slumped sitting [1;2;3]. Position-holding economy can be improved by training [4]. In both mentioned regions, the biomechanical parameters established by myotonometry are significantly affected by the body position [5;6;7]. When force is generated during contraction, the muscle turns more elastic, stiff, and tense [8;9;10;11;12;13]. Vain et al. [14] showed also that the standing effort is not reflected in the elasticity but stiffness of *musculus gastrocnemius*. In the relaxed muscles these characteristics are affected by stretch [12;15;16] and physical fitness is also reflected in these properties [17].

In the “stretch forward flexion” manoeuvre a clearly visible function at low back muscles can be observed in the absence of change in neuromuscular activity [18;19;20]. It has been suggested that behind this phenomenon is the change in the biomechanical properties of the regional passive tissues, which are able to support the upper body without the necessity to increase the neural activation of the muscles [1;18;21;22]. The “sit and reach” manoeuvre is shown to be limited by the tension and elasticity of the thigh muscles [15]. Liu et al. [23], assessed the reliability of this method specifically on trapezius muscle with objective detection of 14.2% increase in stiffness upon shoulder flexion between 0° and 60° and concluded that this is a feasible tool to assess pathological conditions or the effect of treatments.

The pressure in the inter-vertebrate compartments has been measured directly and found to be strongly dependent on the body posture [24]. This must be at least partly a result of the gravity force and partly of structurally formed mechanical levers. This occurs despite the probable absence of surface electromyography (sEMG) activity in regional muscles. At contraction, the pressure in the muscle compartments show a high correlation with muscle tension [9]. The real low-gravity environment achieved in parabola flight or in Space causes specific changes in the biomechanical properties of the muscle [25;26], which are the same as on ground-based low-gravity stimulation [5;6;27;28]. In the Space, the human body takes a gravity-neutral body position where the BTA is around 128° (NASA Man Systems Integration Standards STD-3000, Volume 1, Revision B, 1995). The main reason for this is obviously the balanced tension from antagonistic muscles stabilizing the hip joint without any external interruptive force. The dysplasia of the hip joints in children is healed in a similar position (Pavlik and Bock harnesses; Frejka pillow). According to Harrison et al. [29], already in 1947 Geoffrey McKay Morant recommended for the British Royal Air Force that the pilot’s back-thigh angle (BTA) in the aircraft should be 110–125° for the rest position and 110° for alertness, although this was to be achieved by seat-back inclination. They also claimed in this review article that in the case of the car seats the lowest intra-vertebral disk pressure and regional sEMG activity is inherent to the backrest inclination of 110–130° with concomitant lumbar support. Chen et al. [30] showed that drivers whose BTA angle was less than 86° reported back pain 5 times more often than those with a BTA over 91°.

The aim of the present study was to correlate the muscle’s biomechanical stiffness and the neuromuscular activity characteristics in the shoulder and low-back area in standing, prone, and in different sitting postures. We hypothesized that this parameter might behave separately from the neuromuscular activity of these muscles.

## 2. Methods

### 2.1 Participants

Nine healthy physically active men aged 19–46 years (mean±SD, 28.6±10.9 years), with no complains volunteered for the study. The participants gave their written informed consent prior to the testing. General parameters (mean±SD; minimum-maximum): body mass 78.0±8.2 (66-91) kg, height 182.3±1.57 (180–185) cm, and body mass index (BMI) 25.9±2.8 (20.4-30.0) kg·m^−2^. They were asked to avoid any vigorous physical activity on the previous and on the same day before the study. The protocol of the study was approved by the Ethics Committee of the University of Tartu, nr. 227/T-1.

### 2.2 Muscle stiffness

A MyotonPRO device (Myoton Ltd, Estonia) was used. The device gives a short-lasting (15 ms) mechanical impact (0.4 N) perpendicularly to the tissue with a specific testing end. Oscillation of tissues occurs, and the device follows the dampening of the oscillation with the same testing end. Acceleration of the testing end is monitored at a 3 kHz frequency and a graphic is created (Figure 1) [14;31]. Impact is sufficient to involve muscle tissue [32]. Stiffness reflects the resistance of the tissue to the force that changes its shape and is calculated by the formula C=m·a_1_/Δl(N/m). The higher the value, the stiffer is the tissue, showing that more energy is needed to modify muscle shape.

**Figure 1:**
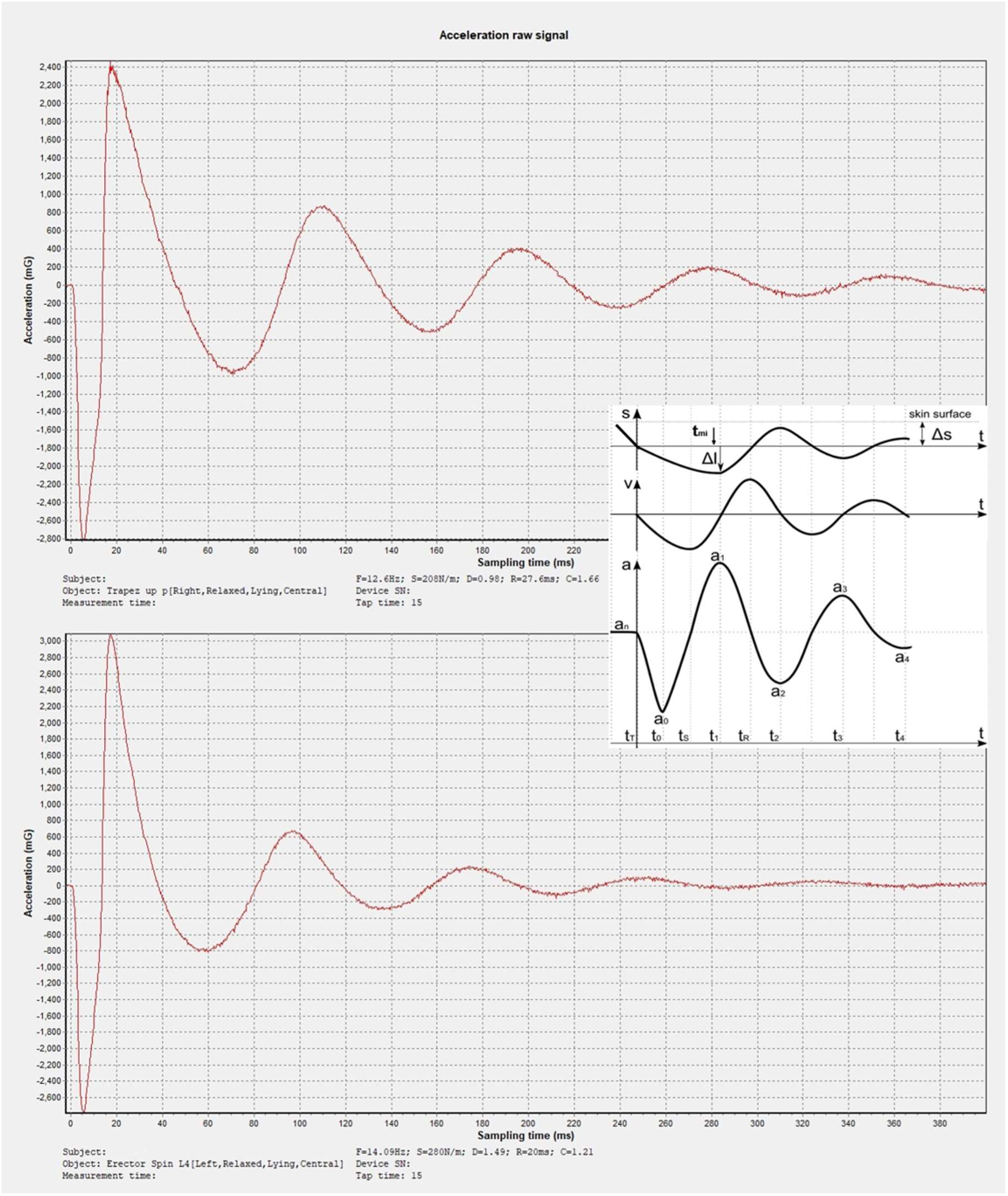
Actual acceleration graphs from measured muscles in lying and schematic graph (adapted from Vain et al. 2015) for calculating stiffness in myotonometry Trapezius muscle in upper and erector spinae in lower graph. Stiffness (N/m) reflects the resistance of tissue to the force that changes its shape; C=m(0.4N)×a_1(max)_/Δl (N/m). This is detected during the initial impact. The higher the value, the stiffer is the tissue, showing that more energy is needed to modify tissue shape.

The measuring point of *musculus trapezius* (upper part, UT) was located and marked on the middle distance between the acromion and C7 vertebra, the latter located by *processus spinosus*. For *musculus erector spinae* (ES), the measuring point was chosen on the level of L4 vertebra located by *spinae iliaca*, and marked in the centre of the muscle belly detected as the most prominent area when extending the torso in the prone position. These regions and muscles were chosen because they are most vulnerable in sitting conditions and they were accessible to the device in the decided body positions.

### 2.3 sEMG activity

The Mean Power Frequency (MPF, Hz) describes the synaptical activation pattern and the Mean Amplitude (AMP, μV) value describes the gross innervation input (the amount of activated motor units) of the selected muscle for a given task [33]. The SENIAM European recommendations for surface electromyography were followed [35]. A telemetric EMG system ME6000 with MegaWin data acquisition software (Mega Electronics Ltd, Finland) was used. Standard bipolar skin electrodes with an inter-electrode distance of 2 cm (Noraxon, USA) were attached to the skin around the mark for the Myoton. Subjects were asked to contract the corresponding muscle groups to ensure that the muscle’s bioelectrical activity can be correctly acquired [36]. The frequency band ranging from 1 Hz to 1 kHz was accepted and the recorded signals were sampled at 1 kHz.

### 2.4 Procedures

A conventional office table (height 75 cm from the floor), to allow the participants to rest their hands on, and a conventional chair (90/90/90°) with a seat height of 48 cm (chair-I) were chosen from the inventory at hand to represent the actual “field/office” conditions. An experimental chair (chair-II) with a convex base and a seat height of 48 cm from the floor was used. This chair allowed the participants to sit up straight while obtaining the back-tight-angle (BTA) 115-120°. The BTA was measured by a standard goniometer (Model 01135, Lafayette Instrument, USA).

Prone, standing, and three sitting postures were adopted and the bilateral measuring was conducted in the following order:

(1) The participants were required to maintain a relaxed standing posture (hands flexed from the elbow at 90° and fingers gently touching the wall to secure their balance, 10 min).
(2) Lying relaxed prone for the measurement on the massage table (10 min).
(3) Sitting straight on chair-I for 5 min (BTA=90°).
(4) Slumping forward and holding for next 5 min (BTA<90°).
(5) Lying on the massage table for 10 min to equalize for the following sitting task.
(6) Sitting on chair-II with a straight back and BTA 115-120° (presented as BTA>90° in table 1 and in figure 2) for 10 min.

**Figure 2.**
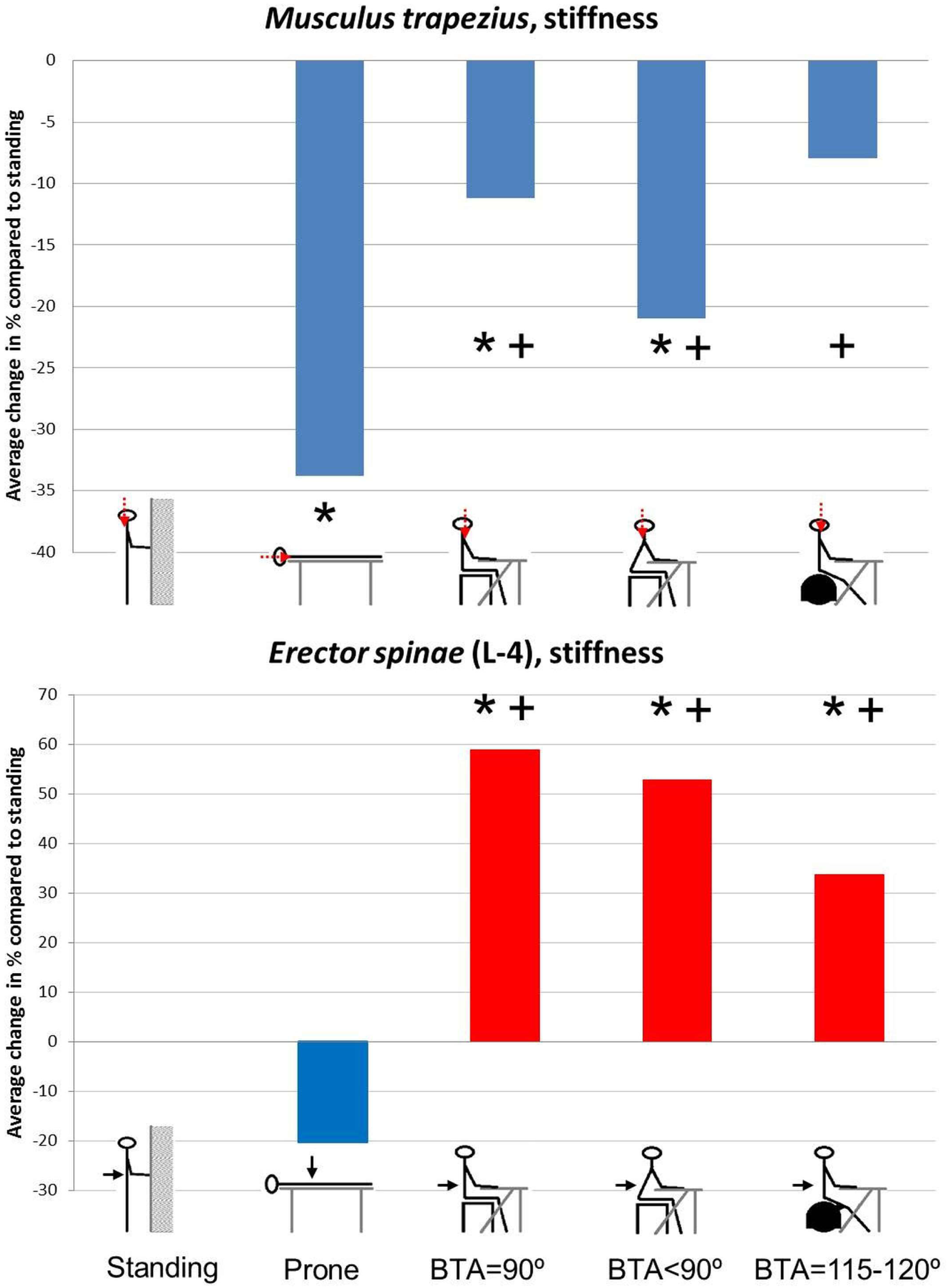
Average differences in stiffness caused by position Average value presenting each Position-group is compared to standing in percentages. Data is presented in table 1. Muscles; upper part of *musculus trapezius* (UT) and *musculus erector spinae* (ES) at the level of L-4. BTA=115 to 120° is presented as BTA>90° in the table 1. * significant difference (p<0.05) compared to standing posture, **+**significant difference to horizontal/prone position.

**Table 1.** Parameters of neuromuscular activity and muscle’s mechanical stiffness.

| Muscle | Parameter | Position | Value, $n=18$ | Lying | Standing | BTA=90 | BTA<90 |
| --- | --- | --- | --- | --- | --- | --- | --- |
| UT | MPF<br>(Hz) | Lying | 65.5±8.0 | * |  |  |  |
|  |  | Standing | 65.9±9.6 | 0.81 | * |  |  |
|  |  | BTA=90 | 69.1±9.6 | 0.16 | 0.30 | * |  |
|  |  | BTA<90 | 64.5±9.6 | 0.84 | 0.81 | 0.24 | * |
|  |  | BTA>90 | 69.0±9.5 | 0.13 | 0.13 | 0.87 | 0.11 |
|  | AMP<br>(μV) | Lying | 3.8±1.2 | * |  |  |  |
|  |  | Standing | 6.1±2.2 | 0.01 | * |  |  |
|  |  | BTA=90 | 5.8±2.1 | 0.02 | 0.40 | * |  |
|  |  | BTA<90 | 4.8±1.1 | 0.05 | 0.12 | 0.49 | * |
|  |  | BTA>90 | 5.7±2.2 | 0.01 | 0.84 | 0.62 | 0.15 |
|  | Stiffness<br>(N/m) | Lying | 181.0±31.7 | * |  |  |  |
|  |  | Standing | 273.5±40.2 | 0.0001 | * |  |  |
|  |  | BTA=90 | 242.9±31.3 | 0.0001 | 0.04 | * |  |
|  |  | BTA<90 | 216.1±33.8 | 0.006 | 0.0002 | 0.03 | * |
|  |  | BTA>90 | 251.7±47.6 | 0.0001 | 0.09 | 0.49 | 0.02 |
| ES | MPF<br>(Hz) | Lying | 61.7±8.0 | * |  |  |  |
|  |  | Standing | 77.6±14.6 | 0.001 | * |  |  |
|  |  | BTA=90 | 71.7±11.9 | 0.01 | 0.31 | * |  |
|  |  | BTA<90 | 68.4±12.5 | 0.28 | 0.06 | 0.40 | * |
|  |  | BTA>90 | 66.7±12.6 | 0.34 | 0.02 | 0.18 | 0.72 |
|  | AMP<br>(μV) | Lying | 3.4±0.8 | * |  |  |  |
|  |  | Standing | 5.4±1.3 | 0.0001 | * |  |  |
|  |  | BTA=90 | 6.0±1.9 | 0.0001 | 0.41 | * |  |
|  |  | BTA<90 | 4.1±1.1 | 0.21 | 0.01 | 0.001 | * |
|  |  | BTA>90 | 4.1±1.0 | 0.05 | 0.01 | 0.001 | 0.70 |
|  | Stiffness<br>(N/m) | Lying | 307.7±65.9 | * |  |  |  |
|  |  | Standing | 386.1±98.7 | 0.07 | * |  |  |
|  |  | BTA=90 | 613.9±134.3 | 0.0001 | 0.0001 | * |  |
|  |  | BTA<90 | 590.1±173.9 | 0.0001 | 0.0001 | 0.57 | * |
|  |  | BTA>90 | 516.6±124.0 | 0.0001 | 0.003 | 0.06 | 0.08 |

This routine was repeated twice to avoid the possible influence of skin-glued electrodes on the close range to the site of myotonometry measuring. First the myotonometry was conducted, the muscles being measured from left to right starting from the UT and then the ES.

### 2.5 Data analysis and statistics

For the analysis of the sEMG results, the data of one-minute recordings in each position were taken and average values were calculated. The time period was concordant to measuring with MyotonPRO in a separate session and this was the first minute of holding each position. For the MyotonPRO device, the MultiScan pattern of 10 continuous measurements with a 1 sec automatic interval was used and the mean result was accepted for future analysis. For finding correlations between neuromuscular activity and the muscle’s biomechanical properties, each muscle was considered separately (calculation: 9 people x 2 sides x 5 positions give *n=*90 for UT; in the case of ES *n*=88 for missing two myotonometric measurements). Differences according to position were calculated with *n=*18 (9 persons x 2 sides) for UT, in case of ES *n=*16 due to the missing data. All values were tested for the normal distribution and only UT MPF showed normal distribution. Therefore, nonparametric statistics were calculated; Spearman ρ to establish correlations, Kruskal-Wallis ANOVA by Ranks, and Mann-Whitney U test were used to compare the averages. The data in the tables are given in means and standard deviations.

## 3. Results

### 3.1 sEMG activity

The age of the participants did not affect any of the measured parameters. However, the effect of individual participants was observed in most of the measured parameters with the exception of the AMP of the UT muscle [Kruskal-Wallis test: H(8, *n=*90)=2.78, p=0.95] and the stiffness of ES muscles [H(8, *n=*88)=11.58 p=0.17]. The position had an effect on all of the measured parameters, except the AMP of UT [H(4, *n=*90)=3.54, p=0.47]. When comparing individual participants, we did not consider the position’s effect.

In ES muscle, the lowest values of motor unit recruitment (AMP) and neural firing activity (MPF) were noted in the lying position. Standing and sitting straight on an ordinary chair (BTA = 90°) showed the most prominent AMP results, differing significantly from the lying and sitting in BTA=115–120° positions (Table 1). Meanwhile, the slumping position (BTA<90°) did not differ from the lying or sitting in BTA=115–120° positions. Concerning the values of neural firing activity (MPF), the standing posture stood solely above lying prone. Interestingly, a statistically significant correlation between both sEMG parameters was observed in the case of the ES muscle [Spearman ρ, *n=*90, ρ=0.24, t(N-2)=2.37, p=0.02] but not in the UT muscle.

### 3.2 Muscle stiffness

In the UT muscle the inter-individual differences were more prominent [H(8, *n=*90)=29.77, p=0.0002]; 16 out of 36 comparisons showed significant differences and 3 other showed p<0.1. Despite the negative Kruskal-Wallis test for the ES muscle [H(8, *n=*88)=11.58; p=0.17], a single individual showed a significant difference from three other respondents (the lowest measured average value 363.6±48.8; *n=*8; N/m, differed significantly from the three highest values 610.2±215.1 N/m, p=0.02; 516.2±103.2 N/m, p=0.002; and 484.4±127.9, p=0.05; *n=*10, 2 regions x 5 positions). The position had a significant effect on both muscles: H(4, *n=*90)=38.91, p=0.00001 in the case of the UT muscle and H(4, *n=*88)=47.86, p=0.00001 in the case of the ES muscle. The values in the lying position were significantly lower from all other positions (Figure 2 depicts group average changes in % compared to standing and without deviation; the exact data is given in Table 1). The measured muscles showed a different response to the change in position. For the UT muscle, almost all positions differed from each other, and support from the table during slumped sitting significantly lowered the values of this parameter compared to straight seated postures. Surprisingly, in the ES muscle there was indifference (p=0.07) between stiffness when standing and lying while both sEMG parameters differed significantly.

### 3.3 Associations between neural activity (sEMG) and biomechanical stiffness

If the whole sample was considered (Table 2), AMP correlated with raised stiffness only in the UT muscle. If each position was considered separately (Figure 3), no significant correlations were observed, although in the standing posture a tendency emerged between stiffness and AMP in the UT muscle (ρ=0.412, p=0.09, *n=*18), also between stiffness and MPF in the ES muscle (ρ=0.438, p=0.07, *n*=18).

**Table 2.**
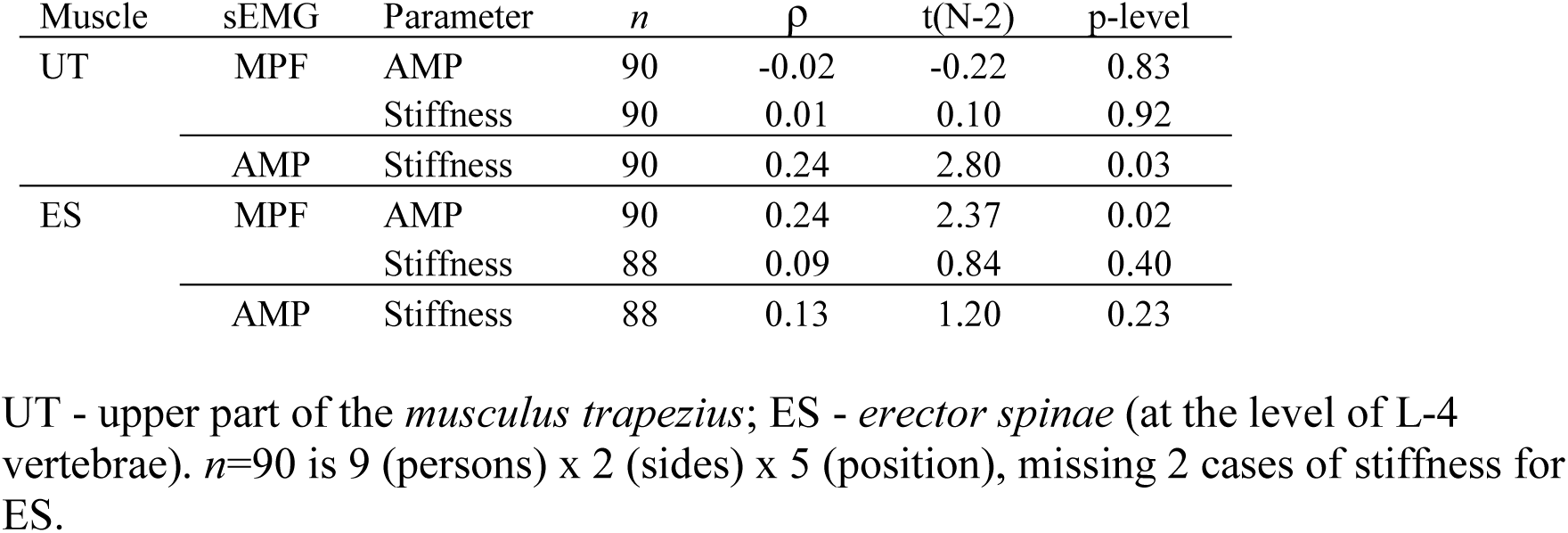
Spearman Rank Order Correlations (ρ)

## 4. Discussion

An increase in sEMG frequency describes a higher moto-neural impulse rate resulting in sustained overlapping of cellular sliding filaments providing higher stiffness transversally. At a higher sEMG frequency, the mechanical impact given transversally (from myoton) to the muscle cells then meets higher resistance from the thicker intracellular environment. We were not able to find in the literature any direct comparisons between neuromuscular and biomechanical characteristics. Concerning the functional state of muscle, in a recent publication stiffness in a biceps brachi muscle was found to be in a positive correlation with time to task failure (TTF), and with the normalized rate of change of mean frequency of sEMG signal [37]. There are also a few studies where this type of comparison was done indirectly [9;13;34]. The latter showed that the resting sEMG amplitudes of the brachial muscles in medicated Parkinson’s disease patients do not differ from the values in the control group, but stiffness does. The authors stated that this is a proof for the affected intrinsic viscoelastic mechanisms in the muscles, which is separated from neural activity. In a recent study Nair et al. [38] used sEMG AMP≤5 μV simply as a criterion to confirm that the muscle is relaxed. Other studies present the results related to the relative contraction force and show that up to 80% of the MVC, the stiffness correlate is almost linear with the generated force [8;11;12].

We consider the most important result of our study that in the measured conditions neuromuscular activity does not correlate with stiffness in the ES muscle. It has been stated previously that lumbar muscle activation depends mainly on the task [39;40;41]. We stress that in our study the BTA 115–120° was not achieved by inclining the back, but by keeping it straight and lowering the thighs when sitting on the chair-II. It seems that this position may be a better alternative to the ordinary chair, although we will not recommend this to ameliorate the well-known harm from prolonged sitting inactivity [42].

We agree with the argumentation that the sEMG silent periods occur due to the strain generated in the elastic connective tissue [1], but wish to emphasize another aspect. Muscle tissue contains perhaps more than 60% of water that by its incompressible nature allows firm support and mediation of force [43]. Another suggestion is that connective tissue contributes actively to stiffness [44]. We can see support for this from ES stiffness results, showing close functional resemblance between the positions of standing and lying during different neuromuscular activities. Although this may be related to the small number of respondents, still, both “natural” positions expressed clearly different stiffness values compared to seated postures while differences were not so prominent in neuromuscular activity.

At contraction, there is a possibility that the correlation between stiffness and neuromuscular activity does exist, for this has been shown indirectly [8]. This allows to explain the findings that the neural activation of the UT muscle must be related to the position of the arms. It also appears that the amount of recruited muscle fibres modifies muscle stiffness more easily than the frequency of neural input (at least in our study). The protein structures in the cells allow important support in the lateral direction [45;46;47]. In line with the above argumentation, we emphasize that input from a single motor neuron is switched to nearby muscle cells in a parallel manner and the separate motor neurons are also switched similarly in parallel, forming specific motor end plate regions [48]. This also seems to allow immediate and firm support in the lateral direction. Perhaps lateral support inside the muscle is a more economical way to ensure arm stability in the muscle, allowing fascia structures to carry the actual load. This may help to explain why in the UT muscle the AMP correlated with stiffness but the firing activity did not. Additionally, in the slumped sitting position (BTA < 90°) the amount of recruited motor neurons is slightly lower than in the other two sitting positions and it shows a tendency to differ from standing (p=0.12). Moreover, the position does not affect the MPF of the UT muscle.

Compared to the lying position, all trunk vertical positions showed significantly higher values in the UT muscle AMP. Leaning forward on the table lowers stiffness on the shoulder area and this may be the main reason for this behaviour in the first place, but this does not ease off the lumbar area for the task centre is transferred there. In this region the obtuse BTA angle instead un-eases regional stiffness in a strong manner and this happens independently from neuromuscular activity.

The neuromuscular activity in the ES muscle did not differ between the standing position and in BTA=90°, showing perhaps an ongoing balancing function. *Musculus iliopsoas* is an important antagonistic muscle supporting spine stability in this region. If antagonistic muscles oppose each other in function then both are neuro-electrically activated. Properties of antagonistic muscles cannot differ in large scale [10], for their synergistic function will be restricted and inefficient [15;49]. Most interestingly, increasing the BTA and keeping the back straight lowers to some extent stiffness in ES compared to both other seated postures and perhaps reflects reaching the optimal angle of the hip joint as in the gravity neutral body position. Steady posture under the Earth’s gravity force seems to be ensured more by biomechanical stiffness than by neuromuscular firing activity. Recruiting the motor units in parallel may allow more effective use of firm support from tissue water and exploit the elastic fascia structures. If this is so, then holding a posture under the constant influence of the Earth’s gravity force becomes a highly economical strategy from such aspects as ATP consuming processes (cross-bridge de-attachment and Ca^2+^ pumping), neurotransmitter management etc. Generation of force is another matter. In that case the force output correlates almost linearly with generated stiffness and tension in the muscles but not with elasticity.

We do consider some of the limitations in our study. One is that the study group is small; however, the changes caused by position and the absence of correlations between stiffness and neuromuscular activity are clear. We did not register the maximal voluntary contraction (MVC) parameters because we were only interested in the actual values of the sEMG results. In addition, the participants were measured around 5pm, and previous physical activity was not recorded. Still, avoiding exhaustive or out of the normal range physical activity during the previous day and prior to the experiment was required and also followed by the participants. On the other hand, this study actually reflects our day-to-day “field” conditions, when we all are influenced by our daily activities, carrying out our sedentary life style.

## 5. Conclusions

The maintaining of the studied steady body positions are to a great extent ensured by biomechanical stiffness of the tissues rather than neuromuscular activity. We also suggest that this is due to the nature of soft tissues, containing mainly incompressible water, allowing a firm mechanical support function. Standing straight and lying, most natural positions for human body, show very similar values in ES stiffness allowing to state that these two positions are the “best for one’s back”. Sitting on an ordinary chair with BTA≤90°, shows the highest stiffness on the low back area, which is in concordance with the general acceptance of the detrimental effect of the sedentary lifestyle. If the seated position is necessary, stiffness in the low back area can be lowered by widening BTA.

## Acknowledgements

This study was partly financed by Enterprise Estonia (EAS, OÜ Ägelind ERDF 25.04.2013). The team would like to express their gratitude to; the inventor of the myotonometry method Mr. Arved Vain PhD, *Dr.Biol.Hab*. for his expertise, and the providers of the experimental chair Mr. Allan Kompus, Mr. Ionel Lehari, and Mr. Toomas Märtson from Ägelind Ltd., To Mr. Viljo Viljasoo who at the time of the study was Associate Professor of the Institute of Technology, Estonian University of Life Sciences. We dedicate this study to the memory of MD PhD Ragnar Viir whose ideas were carrying this research.

## Authors agreement

- Our study was designed to comply with the principles laid down in the Declaration of Helsinki.
- AI and AI-assisted technologies were not used in the writing process.
- Each of the authors has read and concurs with the content in the final manuscript. The material within has not been and will not be submitted for publication elsewhere.
- All authors were fully involved in the; (1) the conception and design of the study, or acquisition of data, or analysis and interpretation of data (2) drafting the article or revising it critically for important intellectual content, (3) final approval of the version to be submitted.
- We confirm that we have given due consideration to the protection of intellectual property associated with this work and that there are no impediments to publication, including the timing of publication, with respect to intellectual property. In so doing we confirm that we have followed the regulations of our institutions concerning intellectual property.
- We understand that the Corresponding Author is the sole contact for the Editorial process (including Editorial Manager and direct communications with the office). He is responsible for communicating with the other authors about progress, submissions of revisions and final approval of proofs.
- We confirm that we have provided a current, correct email or other address which is accessible by the Corresponding Author and which has been configured to accept email from.
- There is no conflict of interest of any sort between the authors and the study presented. This study was partly financed by the Enterprise Estonia (EAS, OÜ Ägelind ERDF 25.04.2013).
- There are no competing interests bound to this study.

